# Regulatory dissection of the severe COVID-19 risk locus introgressed by Neanderthals

**DOI:** 10.1101/2021.06.12.448149

**Authors:** Evelyn Jagoda, Davide Marnetto, Francesco Montinaro, Daniel Richard, Luca Pagani, Terence D. Capellini

## Abstract

Individuals infected with the SARS-CoV-2 virus present with a wide variety of phenotypes ranging from asymptomatic to severe and even lethal outcomes. Past research has revealed a genetic haplotype on chromosome 3 that entered the human population via introgression from Neanderthals as the strongest genetic risk factor for the severe COVID-19 phenotype. However, the specific variants along this introgressed haplotype that contribute to this risk and the biological mechanisms that are involved remain unclear. Here, we assess the variants present on the risk haplotype for their likelihood of driving the severe COVID-19 phenotype. We do this by first exploring their impact on the regulation of genes involved in COVID-19 infection using a variety of population genetics and functional genomics tools. We then perform an locus-specific massively parallel reporter assay to individually assess the regulatory potential of each allele on the haplotype in a multipotent immune-related cell line. We ultimately reduce the set of over 600 linked genetic variants to identify 4 introgressed alleles that are strong functional candidates for driving the association between this locus and severe COVID-19. These variants likely drive the locus’ impact on severity by putatively modulating the regulation of two critical chemokine receptor genes: *CCR1* and *CCR5*. These alleles are ideal targets for future functional investigations into the interaction between host genomics and COVID-19 outcomes.

## Introduction

Since its emergence in late 2019, the SARS-CoV-2 virus has infected more than 160 million people worldwide and claimed more than 3 million lives (WHO, weekly report May 25, 2021). The variance in patient outcomes is extreme, ranging from no ascertainable symptoms in some cases to fatal respiratory failure in others (Vetter et al. 2020). This wide range of patient outcomes is due in part to comorbidities; however, prior health conditions do not explain the full range of outcomes (Zhou et al. 2020). Therefore, efforts have been made to assess a potential genetic component. Repeatedly, a region on chromosome 3 encompassing a cluster of chemokine receptor genes has been reported as having a strong association with an increase in COVID-19 severity in Europeans, with the strongest reported risk variant conferring an odds ratio of 1.88 for requiring hospitalization (p = 2.7*10^-49^, The COVID-19 Host Initiative and Ganna 2021).

Zeberg and Paabo (2020) identified that the COVID-19 severity locus was introgressed by Neanderthals, with a core introgressed haplotype spanning ∼49kb from chr3:45859651-45909024 including rs35044562, reported as one of the leading variants of the association (The COVID-19 Host Initiative 2020) and a broader, extended haplotype with reduced linkage spanning ∼333kb from chr3:45843315-46177096. The core haplotype is at highest frequency in South Asian populations (30%), as well as at appreciable frequency in Europe (8%) and the Americas (4%), yet it is virtually absent in East Asia. The stark difference in frequency between South Asian and East Asian populations implies that the haplotype may have been positively selected in South Asian populations, for which there is support (Racimo et al. 2014; Jagoda et al. 2018; Browning et al. 2018) and/or subject to purifying selection in East Asian populations. However, the specific phenotypic consequences of this haplotype leading to its potential adaptive effect as well as its effect on COVID-19 severity remain unknown. Moreover, the potential causal drivers of this selection, as well as COVID-19 severity remain unstudied.

Here, we identify putative functional variants within this haplotype that may be driving its association with COVID-19 severity. To do so, we first examine the haplotype in the context of a broader introgressed segment. We then identify loci within the introgressed segment that are associated with levels of gene expression (eQTLs) *in vivo*. We next compare the eQTL effects of these variants with differentially expressed genes in COVID-19 and related infection datasets to identify which response genes for these eQTLs are potentially relevant to the COVID-19 phenotype. We follow this computational approach with a high throughput functional Massively Parallel Reporter Assay (MPRA) and identify 20 alleles along the introgressed segment that directly modulate reporter gene expression. We finally intersect these 20 alleles with a host of molecular and phenotypic datasets to further refine them to 4 which display the strongest evidence of contributing to the genetic association with severe COVID-19 at this locus. These shortlisted variants primarily modulate expression through their potential effects on *CCR1* and *CCR5 cis*-regulation and are strong candidate variants that should be investigated with future targeted functional experiments.

## Results

### Genome wide scans for introgression

We carried out two genome-wide searches for introgressed loci in a European population for which we also had available eQTL data using Sprime (Browning et al. 2018) and U and Q95 (Racimo et al. 2014) methods. We used 423 Estonian whole genome samples (Pankratov et al. 2020) that constitute a well-studied representative sample of the broader Estonian population as sampled by the Estonian Biobank (EGCUT) (Leitsalu et al 2015). These samples also have available whole blood RNA-sequence data which contributed to eQTLGen, a broad whole blood eQTL analysis study (Vosa et al. 2018). By utilizing genomes that were part of the eQTL study population, we can be assured that the associations between alleles and gene expression is accurate, as differential linkage disequilibrium (LD) between alleles in different populations can decrease the efficacy of using eQTL data from one population on another.

We initially conducted the Sprime scan (Browning et al. 2018) using the 423 Estonians as the ingroup population along with 36 African samples from the Simons Genome Diversity Project (SGDP) with no evidence of European admixture (Mallick et al. 2016) as an outgroup (Table S1). From this scan, we identified 175,550 likely archaically introgressed alleles across 1,678 segments (Table S2). Following Browning et al. (2018), we then identified segments as confidently introgressed from Neanderthals if they had at least 30 putatively archaically introgressed alleles with a match rate to the Vindija Neanderthal genome (Prüfer et al. 2017) greater than 0.6 and a match rate with the Denisovan genome (Meyer et al. 2012) less than 0.4 (Browning et al. 2018). In total, we identified 693 such segments (Table S3), including the segment containing the COVID-19 severity haplotype on chromosome 3 (see above).

We next used the U and Q95 scan, which specifically identifies regions of introgression showing evidence of positive selection (Racimo et al. 2017). Using Africans from the 1000 genomes project as an outgroup (1000 Genomes Project Consortium, 2015), we found 493 such regions (Table S4). We did not detect the introgressed COVID-19 severity haplotype in our population via this method. This suggests that the COVID-19 severity associated segment, while likely introgressed from Neanderthals based on its detection in our Sprime scan and via the work of others (Zeberg and Paabo 2020), was not under positive selection in the Estonian population. This is consistent with the previously reported lower frequency (8%) of the haplotype in Europeans relative to South Asian populations in which the haplotype is at higher frequency (30%) (Zeberg and Paabo 2020). However, U and Q95 scans do detect this region in South Asian populations (Racimo et al. 2017; Jagoda et al. 2018), supporting positive selection on this haplotype in South Asian, but not European populations.

We next examined which alleles in these putatively Neanderthal introgressed regions detected using these two genome wide scans also are *cis-* and *trans*-eQTLs in the eQTLGen whole blood dataset (Vosa et al. 2018). From the U and Q95 data, we identified 684 *cis*-eQTLs across 250 40kb windows (Table S5). There were no *trans*-eQTLs detected in this set. From the Sprime data, we found 27,342 *cis*-eQTLs from 318 segments along with four *trans*-eQTLs from three segments (Table S6 and S7).

### Refinement of the severe COVID-19 associated introgressed segment

In our Sprime scan, we identified an introgressed region containing the haplotype defined by Zeberg and Paabo (2020) as both introgressed and associated with increased risk of COVID-19-severity. The overall introgressed region as detected in our Estonian population spans ∼811kb from chr3:45843242-46654616, encompasses 16 genes (Figure 1A), and ranks in the top 2% (ranked 21/1677) of Sprime detected segments and in the top 5% (58/1677) of Sprime segments based on length. Its extreme length provides support to the fact that it is introgressed and not likely a product of incomplete lineage sorting, which is detected as seemingly introgressed tracts of significantly shorter length (Huerta-Sanchez et al. 2014).

**Figure 1.**
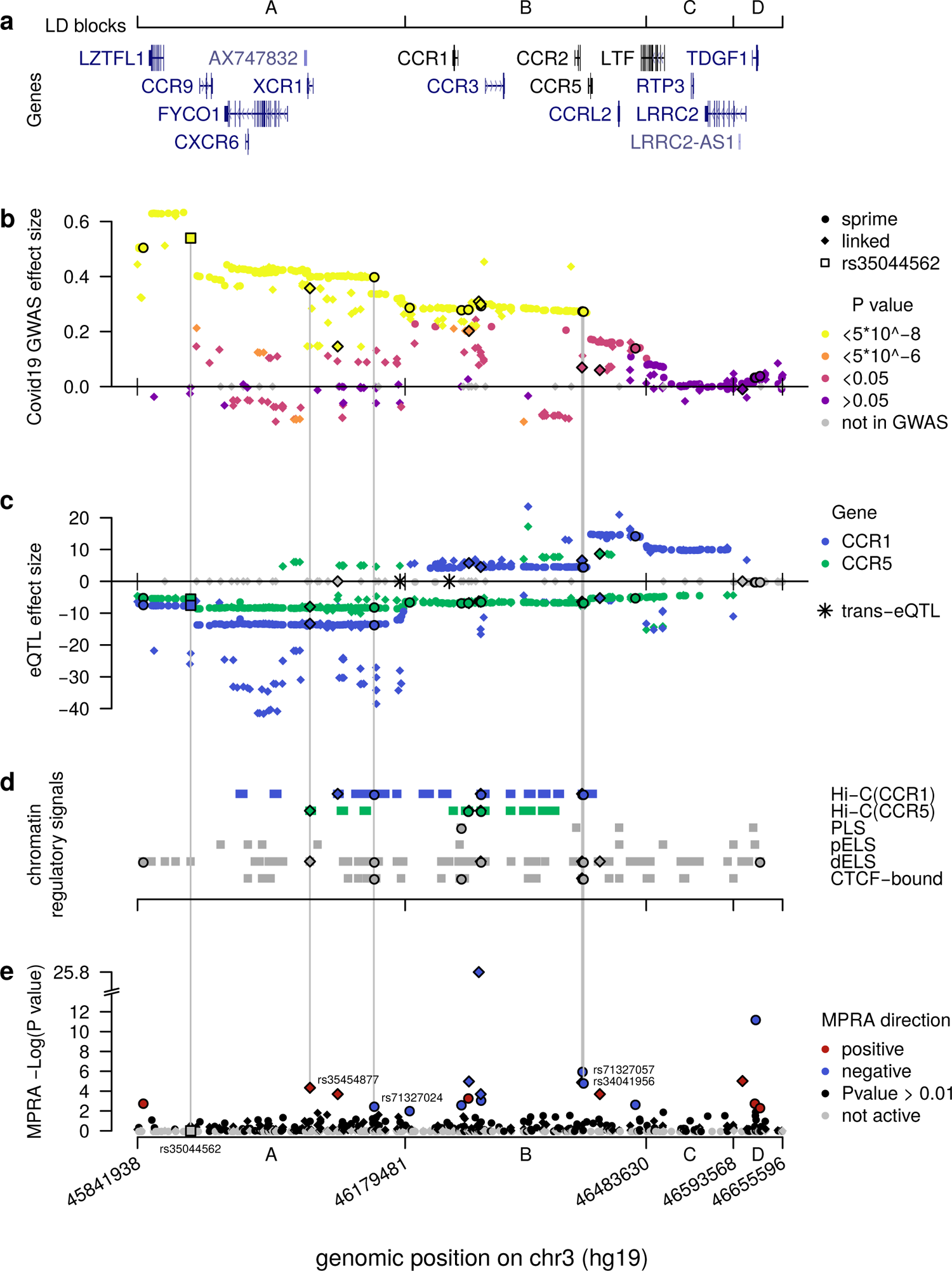
Computational intersections between MPRA emVars and functional genomics datasets across the severe COVIDB19 risk locus. **(a)** Gene locations across the locus along with boundaries of the four LD blocks (ACD). (**b**) Severe COVIDC19 GWAS effect sizes from the release 5 of the COVIDC19 Host Initiative dataset (2021), with strongest genomeCwide pCvalues in yellow spanning the A and B blocks. See key for other color definitions. Dots and diamonds across the panels indicate respectively SNPs identified by Sprime and SNPs in LD with them. (**c**) eQTL effect sizes across the locus (blue for *CCR1*, green for *CCR5*) in whole blood from eQTLGen (Vosa et al. 2018) across the locus. Note the strong downC versus upCregulation of *CCR1* for variants in the A versus B blocks, respectively. Grey SNPs are not eQTLs for any of the two genes or were absent from the eQTL study. Asterisks denote *trans*CeQTLs.(**d**) ChromatinCbased functional annotations across the locus consisting of HiCC contacts with *CCR1* and *CCR5* in Spleen, Thymus or LCL (Jung et al. 2019) and candidate *cis*Cregulatory elements from ENCODE (ENCODE Project Consortium et al. 2020). (**e**) CLog pCvalues for emVars identified using MPRA across the locus. Grey SNPs failed the test for activity in either the archaic or nonCarchaic form. Vertical lines connect the four putative causal emVars and the original tag SNP as described by Zeberg and Paabo (2020) rs35044562 to functional genomics and genetics data. The four putatively causal variants are unique in having significant hits across all functional genomics and genetics tests.

To examine how the introgressed segment may be affecting COVID-19 severity, we began by examining the LD structure within the segment and identified four major blocks defined as minimum pairwise LD between Sprime-identified variants within a block (min r2 = 0.34) (SFig 1). We labeled these blocks as “A” from rs13071258 to rs13068572 (chr3:45843242-46177096), “B” from rs17282391 to rs149588566 (chr3:46179481-46289403), “C” rs71327065 to rs79556692 (chr3:46483630-46585769), and “D” from rs73069984 to rs73075571 (chr3:46593568-46649711) (Figure 1A; SFig 1). All Sprime alleles in the A block are significantly (p < 5*10^-8^) associated with increased risk for COVID-19 severity (The COVID-19 Host Initiative et al. 2021), with the median p-value being 2.32 * 10^-26^ and median effect size being 0.42 (Figure 1B). The B block also harbors many alleles (81.2%) significantly associated with COVID-19 severity, with the median p-value being 1.94*10^-9^ and median effect size being 0.28 (Figure 1B). In the “C” and “D” block no alleles are significantly associated with COVID-19 severity, suggesting that the most likely causal variants for the COVID-19 severity association are found within the A or B blocks (Figure 1B).

All the 361 Sprime-identified introgressed variants act as eQTLs in the whole blood (Vosa et al. 2018) including for many genes that are relevant to COVID-19 infection. Strikingly, of the four *trans*-eQTLs identified genome-wide in our Sprime scan regions in Estonians, two were located on the introgressed COVID-19 severity haplotype. These two variants, rs13063635 and rs13098911, have 11 and 33 response genes, respectively (Table S7). We examined whether these response genes have any relevance to COVID-19 infection and found that 3 (27%) and 13 (39%) of the response genes for rs13063635 and rs13098911, respectively, are differentially expressed in at least one experiment in which a lung related cell-line or tissue was infected with COVID-19 or other related infections (Table S8). These results suggest that these two *trans*-eQTLs may affect the lung response to COVID-19 in a way that could contribute to differential severity in host response. Furthermore, all 361 variants, including the two *trans*-eQTLs, act as *cis*-eQTLs in whole blood, altering the expression of 14 response genes: *CCR1*, *CCR2*, *CCR3*, *CCR5*, *CCR9*, *CCRL2*, *CXCR6*, *FLT1P1*, *LRRC2*, *LZTFL1*, *RP11-24F11.2*, *SACM1L*, *SCAP*, *TMIE*. Of these genes, 7 are chemokine receptor genes (*CCR1*, *CCR2*, *CCR3*, *CCR5*, *CCR9*, *CCRL2*, *CXCR6*), which are likely linked to the segment’s association with COVID-19 severity. There is support in particular for an association between *CCR1* and *CCR5* expression and COVID-19 phenotypes. For example, elevated *CCR1* expression has been demonstrated in neutrophils and macrophages from patients with critical COVID-19 illness (Chua et al. 2020), in biopsied lung tissues from COVID-19 infected patients (Table S8), as well as in Calu3 cells directly infected with COVID-19 (Table S8). *CCR5* expression is also elevated, though to a lesser degree than *CCR1*, in macrophages of patients with critical COVID-19 illness (Chua et al.). Notably, some ligands for *CCR1* and *CCR5* (*CCL15*, *CCL2* and *CCL3*) also show over-expression in these patients (Chua et al.). *CCRL2*, *LZTFL1*, *SCAP*, and *SACM1L* are also differentially expressed in at least one experiment that measures differential expression of genes in lung tissues and related cell lines infected with COVID-19 or other viruses that stimulate similar immune responses (Table S8). These results in general support a role for the introgressed variants along this haplotype contributing to the COVID-19 severe phenotype by affecting the expression of genes that play a role in host COVID-19 response.

Intriguingly, when considering the effect of the *cis*-eQTLs for *CCR1* across the entire segment, we find that the majority of alleles along the introgressed haplotype within the A block are associated with its down-regulation (average Z score = −12.3) (Figure 1C). On the other hand, the majority of alleles within the B and C blocks are associated with *CCR1* up-regulation (average Z scores = 7.1 and 10.2, respectively) (Figure 1C). It is important to note that these eQTL effects are determined based on whole blood from non-infected, healthy patients (Vosa et al. 2018). When considered in the context that severe COVID-19 phenotype is characterized by increased expression of *CCR1* (Chua et al. 2020), these risk-associated alleles having different directions of effect suggest that a complex change to the *CCR1* regulatory landscape driven by alleles across the introgressed segment may be contributing to the disease phenotype. When we consider *CCR5* expression, it shows a more consistent pattern in which the majority of alleles within the A-C blocks are associated with its down-regulation. This result is interesting as *CCR5* expression in patients with severe COVID-19 illness is higher than those with more moderate cases (Chua et al.). However, given the strong LD within each of these segments, discerning the direct connection between one or more alleles driving these regulatory changes and the molecular and phenotypic signatures of severe COVID-19 remains difficult.

### MPRA Variant Selection and Study Design

To independently assess the regulatory impact of the alleles on this COVID-19 risk haplotype, we employed a Massively Parallel Reporter Assay (MPRA) to investigate which alleles on the introgressed haplotype directly affect gene expression. Alleles which have the ability to modulate gene expression in this reporter assay are candidate putatively functional alleles that may drive the association with COVID-19 severity by altering the expression of genes that facilitate the biological response to COVID-19. To ensure that we tested any potential risk variants on the haplotype, we included in the MPRA all variants directly identified in the Sprime scan as being within the introgressed COVID-19 severity associated segment (361), along with any allele linked (r^2^ > 0.3) to one of these Sprime alleles in the Estonian population (140 alleles) or any 1000 Genomes Population (The 1000 Genomes Project Consortium, 2015) European (150 alleles) or South Asian population (197 alleles). Therefore, here we are testing not only alleles on the introgressed haplotype that have a confirmed Neanderthal-specific origin, but also alleles along the introgressed haplotype that were either already present in the human population when the haplotype was introgressed, or arose anew in humans (i.e., human-derived alleles) on the introgressed haplotype following its introgression. After filtering for SNPs falling within simple repeat regions (Benson et al. 1999), which are not compatible with the MPRA (Tewhey et al. 2016), we identified a total of 613 experimental variants. Of these variants, 293 are significantly (p < 5*10^-8^) associated with COVID-19 severity and another 15 approach significance (p < 5*10^-6^), whereas 118 were not tested in the original GWAS (The COVID-19 Host Initiative Release 5) (Figure 1B).

We conducted this assay in K562 cells, a leukemia cell line that displays multipotent hematopoetic biology, which allows for comparison between the MPRA data and the eQTLs identified on whole blood samples (see above). Furthermore, K562 cells can be induced into immune cell fates highly relevant to the COVID-19 severity phenotypes including monocyte, macrophage, and neutrophils (Tabilio et al. 1983; Sutherland et al. 1986; Butler et al. 2014). Moreover, as K562 cells robustly grow and are transfectable using MPRA reagents, they permit the rapid, repeated acquisition of large numbers of cells, as observed in prior MPRAs (Ulirsch et al. 2016; Ernst et al. 2016). Finally, the availability of other published datasets generated on K562 cells (e.g. chromatin ChIP-seq data), allows for comparison between MPRA results, which are episomal in nature, and the endogenous behavior of the genome in the same cell type. However, we do note that MPRA results will also be limited as they will not directly reflect the response of alleles in the endogenous genome and within the *in vivo* tissues in which the COVID-19 response occurs. We therefore also integrated the MPRA results with datasets derived on endogenous immune tissues/cells to help improve our ability to identify biologically relevant candidate driver variants.

### MPRA Reveals 20 Expression Modulating Variants (emVars)

We built the MPRA library following Tehwey and colleagues (2016) and performed four replicates of the experiment in K562 cells (Methods). We observed that normalized transcript counts between replicates were highly correlated (Pearson’s R > 0.999 p = p-value < 2.2e-16) (Figure 2A). As with other MPRA studies (Tewhey et al. 2016; Uebbing et al. 2021), transcript counts in the cDNA samples are significantly correlated with, but not completely explained by sequence representation in the DNA plasmid pool (correlation between means: Pearson’s R = 0.24 p = 2.2e-16; Spearman’s ρ = 0.912 p-value < 2.2e-16) (Figure 2B), suggesting that while some sequences do not have an effect on transcription, other do. The expression of positive control sequences in this experiment was significantly correlated with their expression in the source MPRA (Pearson’s R = 0.59, p = 0.0058) (Figure 2C). Any deviation between positive control sequence activity in this assay and the source MPRA for the two control sets is likely due to additional regulatory information in this assay for which the tested sequences are 270bp compared with 170bp in the source MPRA. Moreover, for our positive control set we observed that 95% of control sequences displayed activity, whereas only 14% of the negative control sequences displayed activity (Figure 2D).

**Figure 2.**
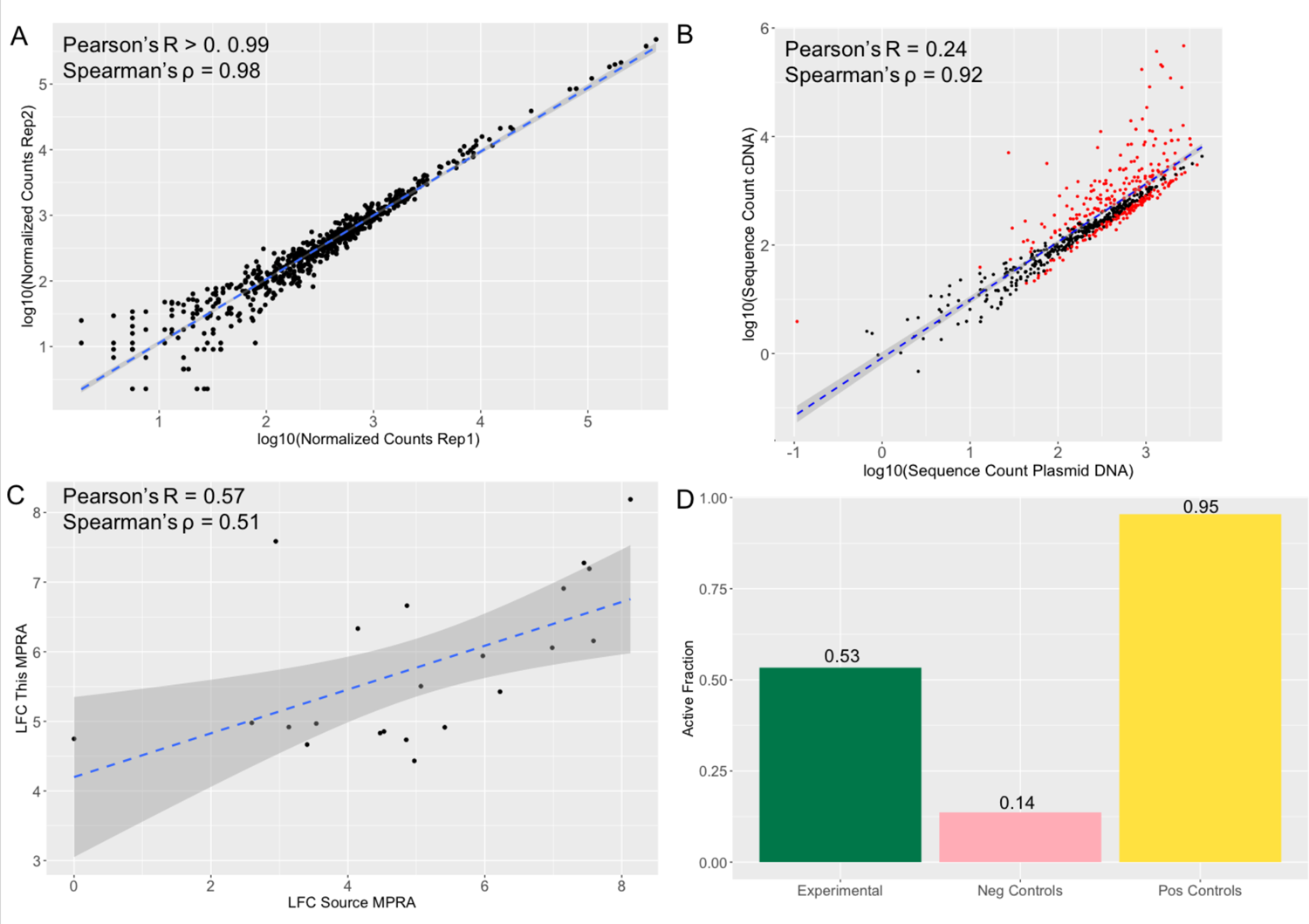
MPRA results show reproducibility and accuracy. **(a)** Log normalized counts for each tested sequence in replicate 1 compared with the replicate 2 of the MPRA. Pearson’s R and Spearman’s ρ are extremely high and significant across pairwise replicate comparisons of all 4 replicates (R > 0.99 pHvalue < 2.2 *10^H16^ M ρ = 0.98 pHvalue < 2.2 *10^H16^). **(b)** Log normalized sequence counts for each tested in the plasmid DNA averaged across the 4 replicates compared with log normalized average sequence counts in the cDNA averaged across the 4 replicates. As with other MPRA studies (Tewhey et al. 2016M Uebbing et al. 2021), there is a significant correlation but the plasmid counts do not fully explain the cDNA counts (Pearson’s R = 0.24 pHvalue < 2.2 *10^H16^M Spearman’s ρ = 0.92 pHvalue < 2.2 *10^H16^), suggesting that some of the sequences have an effect on transcription. Sequencing determined to be significantly active in the MPRA (methods) are colored in red, nonHsignificant points are black. **(c)** Activity (LFC cDNA:pDNA) of positive control sequences in the source MPRA (Jagoda et al. 2021) and in this MPRA. The significant correlation (Pearson’s R = 0.57 pHvalue = 0.006M Spearman’s ρ = 0.51 pHvalue 0.016) suggests that the activity results in this MPRA are accurate. **(d)** Fraction of sequences tested showing significant activity (LFC cDNA:pDNA corrected pHvalue > 0.01). 95% of positive control sequences tested and 0.14% of negative control sequences tested show activity once again suggesting accuracy in the MPRA results. 53% of experimental sequences show significant activity.

Of the 613 experimental variants (1226 alleles) tested, 327 (53%) variants were within sequences found to have detected effects on reporter gene expression (i.e., they are considered “active” or cis-regulatory elements (CREs)) in the context of either the allele on the introgressed haplotype or via its alternative variant (Figure 2D) (Table S9). Consistent with other MPRA studies (Tewhey et al. 2016, Ulirsch et al. 2016; Uebbing et al. 2021), most active sequences showed relatively small effects, with only 17.1% of active sequences showing a LFC greater than 2 (Figure 3A). To confirm that these active CRE sequences reflect endogenous K562 biology, we compared the distribution of active CREs with K562 chromatin state data (Ernst and Kellis 2017; Sloan et al. 2016). We observed that active CREs are significantly enriched relative to non-active sequences for falling with K562 DNase I Hypersensitivity Sites (DHS) and within poised promoters (OR: 8.05 p: 0.023) (Figure 3B). They are also borderline significantly depleted of falling within heterochromatin (OR: 0.73 p: 0.072) (Figure 3B).

**Figure 3.**
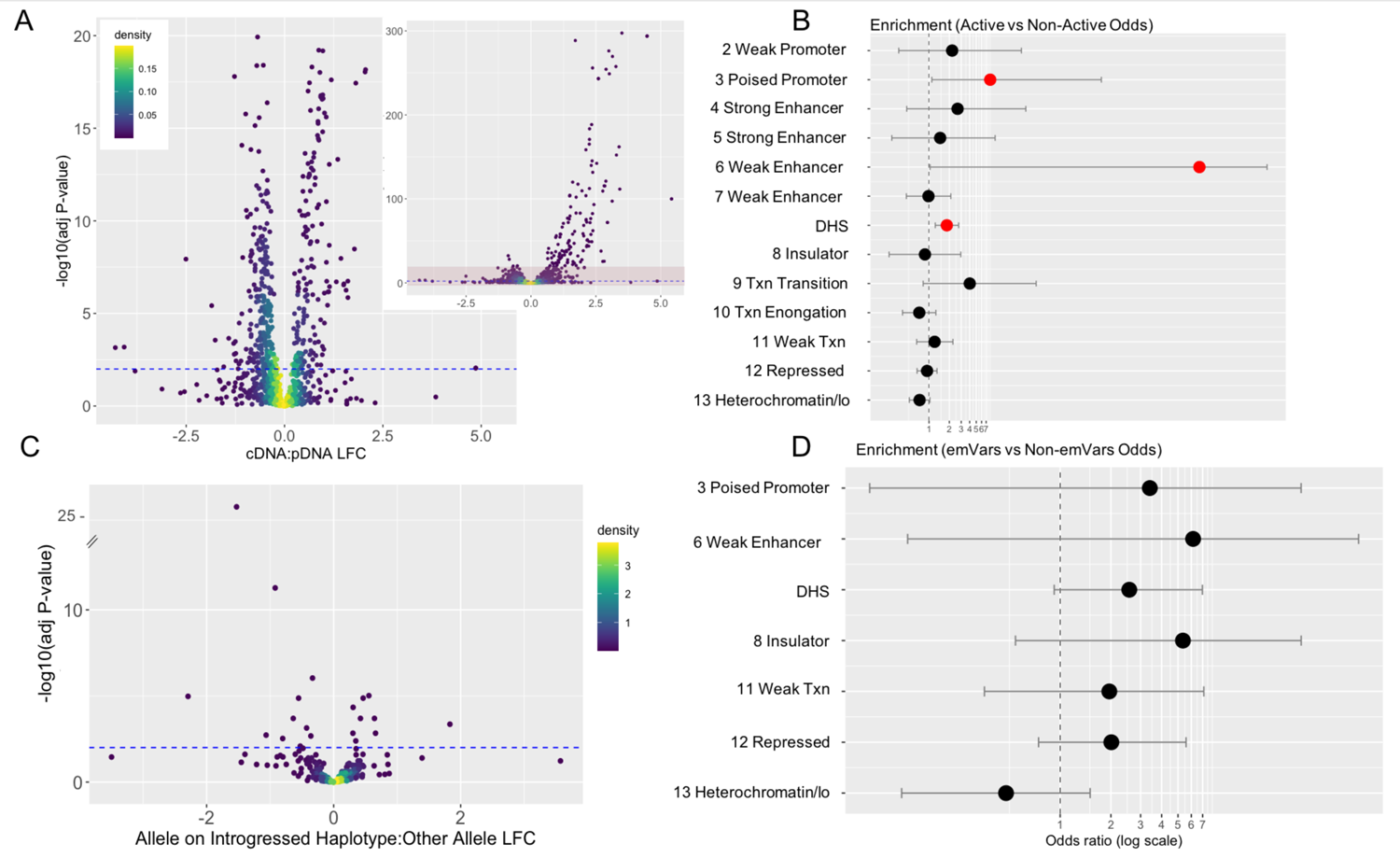
Properties of MPRA3Identified Active Sequences and Expression Modulating Variants. **(a)** Log Fold Change of the cDNA count compared to the plasmid DNA for each sequence and the <log_10_ associated multiple hypothesis corrected p<value. Active sequences are those with a corrected p<value < 0.01D this threshold is denoted with a blue dashed line. The larger plot has a y<axis limit of 20D the inset shows the full spread of the data with the red box denoting the area show in the larger figure. **(b,d)** Enrichment of active sequences in K562 genomic features relative to non<active sequences (b) or sequences with emVars relative to sequences without emVars (d). Genomic features indicated with a number (i.e., “1_”,”2_”) represent chromatin states in K562 cells as defined by Ernst and Kellis (2017). DHS derives from ENCODE (Butler et al., 2014). Enrichments are reported as Fisher’s odds ratios with lines indicating confidence intervals. Significant enrichments (p < 0.05) are colored in red. Missing chromatin states had no overlap with either active sequences (b) or those containing emVars (b). **(c)** Log Fold Change between active sequences with the allele on the introgressed haplotype compared with the sequence with the other allele. Expression modulating variants (emVars) are those whose LFC for this measure is significant with a corrected p<value < 0.01D this threshold is denoted with a blue dashed line.

We next defined as “expression modulating variants” (emVars) those variants exhibiting a significant difference of expression between their two allelic versions using a multiple hypothesis corrected p-value less than 0.01. Using this approach, we identified 20 emVars among the 613 variants we tested (Table 1, Figure 1E, Figure 2B). Consistent with previous MPRA studies (Tewhey et al. 2016, Ulirsch et al. 2016), the effect sizes most emVars detected here are relatively modest, with only 1 emVar having an absolute LFC greater than 2 (Table 1, Figure 3C). Because the sample size is quite small (20), CREs containing emVars do not show significant enrichment within any endogenous K562 functional annotations. However, compared with tested sequences that do not contain emVars, CREs containing these emVars trend towards being over-represented within poised promoters, weak enhancers, DHS, insulator, repressive marks, and for depletion in heterochromatin regions (See Figure 3D).

**Table 1.** (Properties of emVars and Prioritization. This table shows a summary of all the functional data we obtained on the 20 significant emVars identified by the MPRA which we used for prioritization of which of these 20 emVars show the most evidence for contributing to the severe COVIDD19 phenotype. Specifically, we looked for (1) concordance between the eQTL data (Columns H,I) and the HiDC Data (K) D specifically for emVars that are eQTLs for a COVIDD19 relevant gene with which they also physically interact in an immune tissue; (2) a significant association between the allele and severe COVIDD19 in the GWAS data (column J); (3) overlap between the emVar and an Encode annotated cCRE (column L) with support in at least one “class A” immune tissue (column M). The 4 variants that met all these criteria are highlighted in yellow. Greyed out cells show failure to meet a prioritization step. For visualization of this data see Figure 1, for more data from each of these datasets see Table S9.

| snp | chr:pos_hg19 | Introgressed/<br>Allele on<br>Introgressed<br>Hap | Sprime<br>(1) /<br>Linked (0) | LD Block | Distance from<br>rs35044562<br>(kbp) | MPRA<br>Skew<br>Direction | CCR1<br>eQTL<br>Direction | CCR5<br>eQTL<br>Direction | -<br>log10(GW<br>ASp) | HI C Data<br>(thymus, spleen,<br>LCL) | cCREs | cCREs Class A Immune Tissue |
| --- | --- | --- | --- | --- | --- | --- | --- | --- | --- | --- | --- | --- |
| rs17712877 | 3:45848760 | G/C | 1 | A | -60.3 | + | - | - | NA | x | dELS | none |
| rs35454877 | 3:46059484 | T/C | 0 | A | 150.5 | + | - | - | 23.6 | FYCO1,CCR5 | dELS | CD14-positive monocyte female donor ENCDO265AAA |
| rs35657218 | 3:46094462 | A/G | 0 | A | 185.4 | + | 0 | 0 | 8.2 | CCR1,LZTFL1 | x | x |
| rs71327024 | 3:46140073 | G/T | 1 | A | 231 | - | - | - | 24.7 | CCR1 | dELS,CTCF-bound | CD14-positive monocyte female donor ENCDO265AAA, MM.1S, OCI-LY7, GM12878, PC-9, fibroblast of dermis, B cell adult |
| rs13083881 | 3:46184620 | C/G | 1 | B | 275.6 | - | 0 | - | 10.3 | x | x | x |
| rs34919616 | 3:46250008 | G/A | 1 | B | 341 | - | 0 | - | 9.5 | x | PLS,CTCF-bound | CD14-positive monocyte female donor ENCDO265AAA, MM.1S, GM12878 |
| rs35617677 | 3:46258740 | C/T | 1 | B | 349.7 | + | 0 | - | NA | CCRL2,CCR5 | x | x |
| rs34985947 | 3:46260008 | C/T | 0 | B | 351 | - | + | - | 6.6 | CCRL2,CCR5,CCR2 | x | x |
| rs1542755 | 3:46272440 | G/T | 0 | B | 363.4 | - | 0 | - | 10.8 | CCR2 | x | x |
| rs13096905 | 3:46274766 | G/A | 1 | B | 365.7 | - | 0 | - | 9.3 | CCR5,CCR1,CCRL2,CCR2,CCR2 | dELS | none |
| rs10510750 | 3:46274886 | T/C | 0 | B | 365.9 | - | + | - | 9.5 | CCR5,CCR1,CCRL2,CCR2 | dELS | none |
| rs762789 | 3:46402627 | G/A | 0 | B | 493.6 | + | + | - | 1.6 | CXCR6,CCR1 | dELS,CTCF-bound | CD14-positive monocyte female donor ENCDO265AAA |
| rs71327057 | 3:46402645 | A/C | 1 | B | 493.6 | - | + | - | 7.9 | CXCR6,CCR1 | dELS,CTCF-bound | CD14-positive monocyte female donor ENCDO265AAA |
| rs34041956 | 3:46402734 | G/A | 1 | B | 493.7 | - | + | - | 7.8 | CXCR6,CCR1 | dELS,CTCF-bound | CD14-positive monocyte female donor ENCDO265AAA |
| rs6777875 | 3:46425092 | G/T | 0 | B | 516.1 | + | - | + | NA | x | dELS | B cell adult, MM.1S, OCI-LY7 |
| rs34245951 | 3:46469384 | T/C | 1 | B | 560.4 | - | + | - | 2.7 | LZTFL1 | x | x |
| rs73072002 | 3:46605033 | C/T | 0 | D | 696 | + | 0 | 0 | 0.1 | ALS2CL,LTF | x | x |
| rs55649212 | 3:46620020 | A/G | 1 | D | 711 | + | 0 | 0 | 0.3 | x | x | x |
| rs2293021 | 3:46621117 | A/G | 1 | D | 712.1 | - | 0 | 0 | 0.3 | x | x | x |
| rs73073752 | 3:46626599 | C/T | 1 | D | 717.6 | + | 0 | 0 | 0.3 | x | dELS | none |

### Evaluation of emVars with Other Functional Data Reveals Four Putatively Causal Variants

We next examined the 20 emVars for additional evidence of a role in the regulatory mechanisms potentially linked with COVID-19 response. Particularly, we looked for emVars that are (1) significantly (p < 5*10^-8^) associated with COVID-19 severity, (2) are eQTLs for a gene with strong evidence of relating to the severe COVID-19 phenotype, (3) are within a chromosomal region that that based on Hi-C data physically interacts with the promoter of the COVID-19 associated gene in an immune cell line, (4) are within chromatin regions that have with epigenomic marks consistent with acting as *cis*-regulatory elements in human immune cells (see “emVar prioritization” section of Methods for full details) Of the 20 emVars, we found four that meet all of these criteria (Table 1, Figure 1). Based on the combination of eQTL and Hi-C data, the variant rs35454877 is most likely implicated in the down-regulation of *CCR5,* while variants rs71327024, rs71327057 and rs34041956 appear to be involved in the regulation of *CCR1*. Of these, rs35454877 and rs71327024 fall within the LD Block A, which shows the strongest GWAS associations for COVID-19 severity and are 150kb and 230kb downstream of rs35044562, the originally defined tag SNP for this GWAS signal (Zeberg and Paabo 2020). Furthermore, data from other GWAS studies shows that these four variants are significantly (p < 5*10^-8^) associated with other phenotypes that could relate to the COVID-19 severity phenotype including: “Monocyte count”, “Granulocyte percentage of myeloid white cells”, and “Monocyte percentage of white cells” (Astle et al. 2016).

## Discussion

Following our research approach, we re-examined a previously identified adaptively introgressed segment in Eurasians within the context of the genome-wide signatures of introgression. We identified 613 variants within the introgressed region (chr3:45843242-46654616) and these were tested for whether they are potential drivers of the association this region exhibits with increased COVID-19 severity. Using MPRA, we tested these variants in a multipotent immune-related cancer cell line and narrowed down the list to 20 emVars where the expression level driven by the allele on the introgressed haplotype was significantly different from the expression driven by the other allele. We did not find support in the MPRA for the expression modulation potential of rs35044562, the variant previously reported as the tagging GWAS signal for COVID-19 severity (Zeberg and Paabo 2020). This variant may therefore be likely only a tagging rather than a causative variant, albeit functional experimentation on other cell lines may show otherwise. Further mining our MPRA results in concert with datasets on the epigenomic and transcriptional environment of immune cells from other functional genomics sources, here we highlight four emVars that have particularly strong evidence of acting as putatively causal variants and whose archaic alleles are strongly implicated with *CCR1* (rs71327024, rs71327057, rs34041956) and *CCR5* regulation (rs35454877).

We caution that while our experimental design was optimized for detecting *cis*-eQTLs variants effects and within a multi-potent immune-related cancer cell line, other, longer range interactions between genomic regions and in other cell types may also be mediating severe COVID-19 response. For example, when comparing the response genes of the two identified *trans*-eQTLs in the introgressed haplotype to RNA-seq studies testing COVID-19 infection in lung cell lines and tissues (Table S8), 5 (45%) and 14 (41.2%) of the response genes for the two *trans*-eQTLs in this locus rs13063635 and rs13098911, respectively, were detected as differentially expressed in at least one *in-vitro* experiment, although neither of these variants were detected as emVars in this MPRA. Therefore, we also urge additional functional studies to consider the effects of these *trans-*eQTLs and, in general, to replicate our findings in lung epithelial and other cell types from which some of the expression evidence we here build on are derived.

While our study provides strong functional support for at least four archaic variants at the introgressed locus, the direction of the effect of these alleles in both the healthy and infected state needs further clarification. For example, in the severe COVID-19 phenotype, *CCR1* and *CCR5* are upregulated relative to their expression levels in moderate cases of the disease (Chua et al. 2020), while in our MPRA experiments the three top candidate emVars acting as regulators of *CCR1* all showed down-regulation. Furthermore, alleles in Block A of the introgressed haplotype, which exhibit the strongest GWAS associations with COVID-19 severity, all act as down-regulating eQTLs for *CCR1* in healthy whole blood samples. We hypothesize that this difference between the direction of effect of the alleles in the healthy *in vivo* and reporter assay *in vitro* episomal condition relative to in the disease state reflects the fact that these alleles contribute to the risk of severe COVID-19 by destabilizing the regulatory mechanism of *CCR1* and *CCR5,* such that they have decreased expression in the healthy form, but are hyper-expressed upon infection. Additional work needs to be done to further explore this potential mechanism as well as to uncover what therapeutic roots may be undertaken to mitigate this mechanism, should it be found to be accurate.

## Supporting information

SFig1

## Acknowledgements

We would like to thank Drs. Steve Reilly, Maryellen Ruvolo, David Pilbeam, Dan Lieberman, and David Reich, as well as members of the Capellini and Sabeti labs for their critical insight into this work. D.M. and L.P. are supported by the European Regional Development Fund, projects No. 2014-2020.4.01.16-0024, MOBTT53. FM is supported by the European Regional Development Fund, projects No project no. 2014-2020.4.01.16-0030. E.J is supported NSF DDRIG (BCS-1847287) and T.D.C by NSF (BCS-2020205) and The American School of Prehistoric Research, Harvard University.

## Author Contributions

E.J. conceptualized the project, performed the MPRA experiment and the functional follow up analyses, ran the population genetics analyses and was the primary author of the manuscript text. D.M. performed downstream functional analyses, provided interpretation of results and participated in manuscript writing and editing. F.M. assembled the eQTLs dataset, provided interpretation of results, revised the manuscript. D.R. processed external promoter-capture datasets, and performed different-expression analyses for COVID-19 and related RNA-seq datasets, L.P. ran the population genetics analyses, provided interpretation of results and participated in manuscript editing. T.D.C. conceptualized the project, provided supervision for all experiments and analyses, and participated in manuscript writing and editing.

## Data and Code Availability

The data supporting the findings of this study, as presented in all figures (Figures 1-3), including Supplementary Figures (SFig 1), are available within Tables S1-9 included in this publication as well as from the corresponding author upon reasonable request. We have deposited all MPRA data in Geo Omnibus (GSE176233). The following secure token has been created for Reviewers to access the raw datasets: XXXX. eQTL data was obtained from the public repository of the eQTLGen consortium: http://www.eqtlgen.org/cis-eqtls.html, GWAS data was obtained from the COVID-19 Host Genetics Initiative: https://www.covid19hg.org/results/. We used the most recent release (release 5). filename: COVID19_HGI_A2_ALL_leave_23andme_20210107.b37.txt.gz

## Methods

### Introgression Scans

#### SPrime

We downloaded the Sprime software from https://faculty.washington.edu/browning/sprime.jar and ran it using java-v1.8.0_40. We used 423 Estonians as the ingroup population along with 36 African samples from the Simons Genome Diversity Project (SGDP) with no evidence of European admixture (Mallick et al. 2016) as an outgroup (Table S1). Following Browning et al. (2018), we then summed the number of Sprime alleles per segment, and for segments greater than or equal to 30 alleles, we calculated the match rate to the Vindija Neanderthal genome (Prüfer et al. 2017) and the Denisovan genome (Meyer et al. 2012). Segments with greater than 0.6 match rate to the Neanderthal genome and less than 0.4 match rate to the Denisovan genome, were considered introgressed by Neanderthals. In total, we identified 693 such segments (Table S3), including the segment containing the COVID-19 severity haplotype on chromosome 3 (see above).

#### U and Q95

Concerning U and Q95, following Racimo et al. (2017), for every 40kb window within the genome, we calculated the U score as the number of SNPs per 40kb window which had <1% frequency in a combined panel of African (AFR) individuals from the 1000 Genomes project (The 1000 Genomes Project Consortium 2015) that show no major evidence of European admixture (Table S1), had >20% frequency in 423 Estonians, and are homozygous in the Vindija Neanderthal genome (Prufer et al. 2017). We also calculated the Q95 score as the 95% quantile frequency of the derived alleles in the Estonian set that are homozygous for the Vindija Neanderthal allele and are at <1% in the African set. We finally determined the top scoring windows to be those that are in the top 99th percentile of windows in terms of both U and Q95 scores (FDR 0-5.5%) Racimo et al. 2017.

#### GWAS data

GWAS data was downloaded from the most recent release (release 5) from the COVID-19 Host Genetics Institute (The COVID-19 Host Genetics Initiative, Ganna A 2021). We utilized the “A2_ALL_leave_23andme” dataset for which the tested phenotype is “Very severe respiratory confirmed covid vs. population.” There are 5,582 cases and 709,010 controls in this dataset.

#### eQTL data

eQTL data was obtained from the public repository of the eQTLGen consortium http://www.eqtlgen.org/cis-eqtls.html. Only significant *cis-* and *trans-*eQTLs, multiple hypothesis corrected p-value < 0.05 were included. Whenever Z-scores are reported including in Table S5 and S6, the scores are polarized to the correct direction of effect of the allele along the introgressed haplotype.

### Covid-19 and related RNA-seq datasets

Datasets for RNA-seq studies performed on *in-vitro* lung cell lines exposed to either COV-2 infection, related coronaviruses (e.g. MERS), other virus infection (e.g. RSV), or immune stimulation were obtained from the GEO database. Namely, GSE147507 provided by the tenOever lab (Blanco-Melo et al. 2020; Daemen et al. 20201) – Series 1-9 and 15, GSE139516 (Zhang et al. 2020), GSE122876 (Yuan et al. 2019) and GSE151513 (Banerjee et al. 2021). Raw RNA-seq data were all processed with a similar pipeline. Sequence read quality was checked with FastQC (https://www.bioinformatics.babraham.ac.uk/projects/fastqc/), with reads subsequently aligned to the human reference transcriptome (GRCh37.67) obtained from the ENSEMBL database (Hunt et al. 2018) which was indexed using the ‘index’ function of Salmon (version 0.14.0) (Patro et al. 2017) with a k-mer size of 31. Salmon alignment was performed with the ‘quant’ function with the following parameters: “-l A --numBootstraps 100 --gcBias -- validateMappings”. All other parameters were left to defaults. Resulting quantification files were loaded into R (version 3.6.1) (R Core Team 2017) via the tximport library (version 1.14.0) (Soneson et al. 2015) with the “type” option set to “‘salmon”. Transcript quantifications were summarized to the gene level using the corresponding hg19 transcriptome GTF file mappings obtained from ENSEMBL. Count data were subsequently loaded into DESeq2 (version 1.26.0) (Love et al. 2014) using the “DESeqDataSetFromTximport” function. For subsequent differential-expression analysis, a low-count filter was applied prior to library normalization, wherein a gene must have had a count greater than five in at least three samples (in a given dataset) in order to be retained. For tenOever datasets, differential expression analysis was performed comparing treated samples (i.e. infected of stimulated cells) relative to the respective series’ mock control samples. For the Yuan et al. 2019 dataset expression was compared between MERS-infected and mock controls. For the Zhang et al. 2020 dataset, hours post-infection were used as a continuous variable (with mock representing ‘0H post-infection’) for the DESEQ2 model, with significance defined as a gene being up- or down-regulated as a function of post-infection time. The differential expression analysis for the processed Banerjee et al. 2021 dataset, which is also a time-course dataset, was implemented as described previously (Banerjee et al. 2021). Sets of significant genes in each dataset (defined as having a Benjamini-Hochberg adjusted p-value of < 0.05) were subsequently intersected with the sets of response genes identified for the *cis-* and *trans*-eQTLs described in this study.

### MPRA Design and Implementation

We used a MPRA to determine which of these 613 variants within the introgressed segment fall within active *cis*-regulatory elements (CREs) and whether they modulate reporter gene expression relative to the other variant at the same position. We conducted this assay in K562 cells, a leukemia cell line that displays multipotent hematopoetic biology and which allows for comparison between MPRA data and eQTL datasets derived from whole blood samples. Furthermore, K562 cells can be induced into cell fates associated with the COVID-19 phenotypes including monocyte and macrophage and neutrophils (Butler et al. 2014).

Each variant was tested in the context of 270bp of the endogenous sequence centered around each variant. For the SPrime alleles, if there is another allele within the span of this 270bp sequence that is highly linked (r2 > 0.8) in the Estonian population and at least in 9 of the 10 1KG populations it was included in the 270bp sequence. This 270bp sequence will be hereafter referred to as the “tested sequence”. Additionally, we included 44 control sequences from a past MPRA experiment performed in K562 cells (Jagoda et al. 2021) with the 22 strongest up-regulating sequences from this MPRA serving as positive controls and the 22 sequences with smallest magnitude of effect on expression serving as negative controls. In the original assay, these control sequences were 170bp in length, here we extended the sequences to create 270bp sequences centered around the original 170bp sequence. This difference in length could account for any potential regulatory discrepancy between the original experiment and this one. In total, the MPRA experiment consisted of 1,270 sequences. All tested sequences additionally included 15bp of adaptor sequence on both the 5’ and 3’ side to facilitate cloning into the MPRA vector. Following Tewhey et al. 2016, sequences were cloned into the MPRA vectors oriented according with the nearest transcription start site.

The tested sequences were synthesized by Twist Bioscience and the cloning steps to generate the MPRA vector library were conducted following the procedure outlined by Tewhey and colleagues (2016) using the scale of their smaller library size. Barcoded sequences were initially cloned into pGL4:23:ΔxbaΔluc vectors and 4 sequencing libraries were prepared to sequence across the oligos and barcodes to determine oligo-barcode combinations within this mpraΔorf pool. Sequencing was conducted by the Harvard Bauer Core facility on an Illumina NovaSeq using 2 x 250bp chemistry. Again, following Tewhey and colleges (2016), an amplicon containing a GFP open reading frame, minimal promoter and partial 3’ UTR was cloned into the mpraΔorf library to make the final mpra:gfp library.

For each of four biological replicates, 40ug of mpra:gfp vector pool was then transfected into 10 million K562 cells using electroporation with the Lonza 4D-Nucleofector following the manufacturer’s protocol. After 24hrs, cells were collected and flash frozen in liquid N_2_. Closely following Tewhey and colleagues (2016) procedure for their smaller library, total RNA was extracted, GFP mRNA was isolated, converted to cDNA and prepared into sequencing libraries to sequence the barcodes. 4 sequencing libraries were also prepared of the mpra:gfp plasmid to obtain the representation of each barcode in the transfected vector pool. Barcode sequencing was performed on an Illumina MiSeq with 1 x 50bp chemistry at the Harvard Bauer Core.

### MPRA Data Analysis

#### Barcode - Oligo reconstruction

All MPRA data analysis steps were conducted following Tewhey and colleagues (2016). The 250bp paired end reads from the sequencing of the mpraΔorf library were merged using Flash v.1.2.11 (Magoč and Salzberg, 2011). Merged amplicon sequences were then filtered for quality control such that sequences were kept if (1) there was a perfect match of 10bp on the left or right side of the barcode, (2) the 10bp on both sides of the barcode matched with levenshtein distance of 3 or less, and (3) the 2bp on either side of the barcode matched perfectly. Sequences that passed through these filters were aligned back to the expected sequence pool using Bowtie2 v. 2.3.4.1 with the --very-sensitive flag. Alignments that had less than 95% perfect matching with the expected sequence and any alignment which had a mismatch at the variant position were removed. Barcodes that matched to more than one expected sequence are unusable and therefore were also removed.

Because of our small library size, we were able to get a very high barcode yield. All oligo sequences were represented and tagged with a wide diversity of barcodes, with the median unique barcodes per oligo being 18,812 (Figure S2A). Only 5 oligos had fewer than 100 unique barcodes (Figure S2B), with the fewest being tagged by 27 unique barcodes. This large number of unique barcodes lead to extremely high reducibility between replicates (Figure 2A), which will allow for a high degree of sensitivity to detect subtle differences between alleles (Tewhey et al. 2016).

### Tag Sequencing

Again, following Tewhey and colleagues (2016), the 1 x 50bp tag sequencing reads were filtered such that reads were only kept if they had a maximum levenshtein distance of 4 with the constant sequence within the 3’ UTR of the GFP as well as a perfect match with the two base pairs adjacent to the barcode. If the sequence passed through these filters, the barcodes were then matched back to the oligos based on the information from the mpraΔorf library sequencing described above. The counts for each barcode were summed for each oligo. This summation of the counts per barcode reduces the noise that could be derived from any individual barcode having a functional effect.

### Determination of Active Putative *Cis-*Regulatory Elements and Expression Modulating Variants (emVars)

Following Tewhey and colleagues (2016), the summed oligo counts from the tag sequencing for all 4 cDNA samples and all 4 plasmid samples were passed in Deseq2 and sequencing depth was normalized using the median-of-rations method (Love et al. 2014). We then used Deseq2 to model the normalized read counts for each oligo as a negative binomial distribution (NB). Deseq2 then estimates the variance for each NB by pooling all oligo counts across all the samples and modeling the relationship between oligo counts and the observed dispersion across all the data. It then estimates the dispersion for each individual oligo by taking this observed relationship across all the data as a prior a performing a maximum posteriori estimate of the dispersion for each oligo. Therefore, the bias for the dispersion estimate for each oligo is greatly reduced because it relies on pooled information from all other oligos. We then used Deseq2 to estimate whether an oligo sequence had an effect on transcription by calculating the log fold change (LFC) between the oligo count in the cDNA replicates compared with its count in the plasmid pool. We tested whether this LFC constituted a significant difference of expression using Wald’s test and required a stringent Bonferroni corrected p-value of less than 0.01 for a significant result. If an oligo sequence had a significant LFC with either allele, the sequence is considered “active”. Finally, to determine which variants are expression modulating, for oligos which were determined to be active, we used Deseq2 to calculate the fold change between the two versions of the oligo sequences with Wald’s test to calculate the p-values. p-values were then corrected using the Benjamini-Hochberg test to correct for multiple hypothesis testing. Significance was defined stringently as a multiple hypothesis corrected p-value of <0.01.

### emVar Prioritization and intersection of data from other functional sources

To identify which of the 20 variants identified by the MPRA as emVars are the best candidates for contributing to an increased risk of severe COVID-19, we the MPRA data with data from other sources specifically:

### GWAS data

As described above, GWAS data is from the most recent release (release 5) from the COVID-19 Host Genetics Institute (The COVID-19 Host Genetics Initiative, Ganna A 2021). If the p-value for an emVar was less than 5 * 10^-8^ (or -log10(p-value) > 7.3), it was considered significant and passed through this prioritization step.

### Promoter-capture Hi-C datasets

Promoter-capture (Hi-C) data were obtained from Jung et al., 2019, particularly the file ‘GSE86189_all_interaction.po.txt.gz’ containing processed information on genomic regions with significant contacts of targeted promoters. This dataset was generated from promoter-capture assays across a number of different tissues and cell-types; given our particular interest in immune cell regulation, we considered only those significant interactions (reported p-value < 0.01) present in samples from lymphoblasts (GM12878.ADS and GM19240.ADS), spleen (STL001.SX1 and STL003.SX3) and thymus (STL001.TH1) samples. Interacting regions, which may indicate putative cis-regulatory elements, were intersected with our defined emVars using bedtools intersect. If an emVar falls within a contact site for any gene, this is reported in Table (1). For our prioritization, if an emVar is both within a contact site for a gene with relevance to COVID-19 infection, particularly *CCR1,* which is differentially expressed in some SARS-COV-2 infection studies analyzed here (Table S8) and by others (Chua et al. 2020), and *CCR5* which reported as is differentially expressed in other studies (Chua et al. 2020), and the emVar is also an eQTL for this same gene (Vosa et al. 2018), that supports its prioritization (Figure 1, Table 1).

### Candidate cis-Regulatory Elements (CCREs) by Encode

To further validate emVars for biological relevance, we downloaded all 926,535 human cCREs from https://screen.encodeproject.org/ (ENCODE Project Consortium, 2020). cCREs are DNAse hypersensitivity sites that are further supported by additional evidence of *cis*-regulatory activity in the form of either histone modifications (H3K4me3 and H3K27ac) or CTCF-binding data. From there, these cCREs are further classified based on the combination of both their epigenomic signals as well as their genomic context into 4 major categories: cCREs with promoter-like signatures (cCRE-PLS), cCREs with enhancer-like signatures (cCRE-ELS) (these are further subsetted as either proximal (pELS) or distal (dELS), DNase-H3K4me3 cCREs, and CTCF-only.

To intersect these human cCREs with our emVar data, we first used LiftOver (Hinrichs et al. 2006) to convert the our emVar coordinates from *GRCh37 to GRCh38* and then used Bedtools intersect to search for emVars falling within cCREs. These intersections are reported in Table 1. For emVars that already passed through the prior prioritization steps that overlapped a cCRE, we then examined the tissue level information on the cCRE on the web browser (https://screen.encodeproject.org/). emVars were prioritized if they are within cCREs that had cCRE annotations in a cell at least one immune-related “class a” cell line, which is a cell line for which data on all four makers (DNAse, H3K4me3, H3K27ac, CTCF-binding) is available. These results are displayed in Figure 1D and Table 1.

