## Supplementary material for "Regulatory dissection of the severe COVID-19 risk locus introgressed by Neanderthals": SFig1

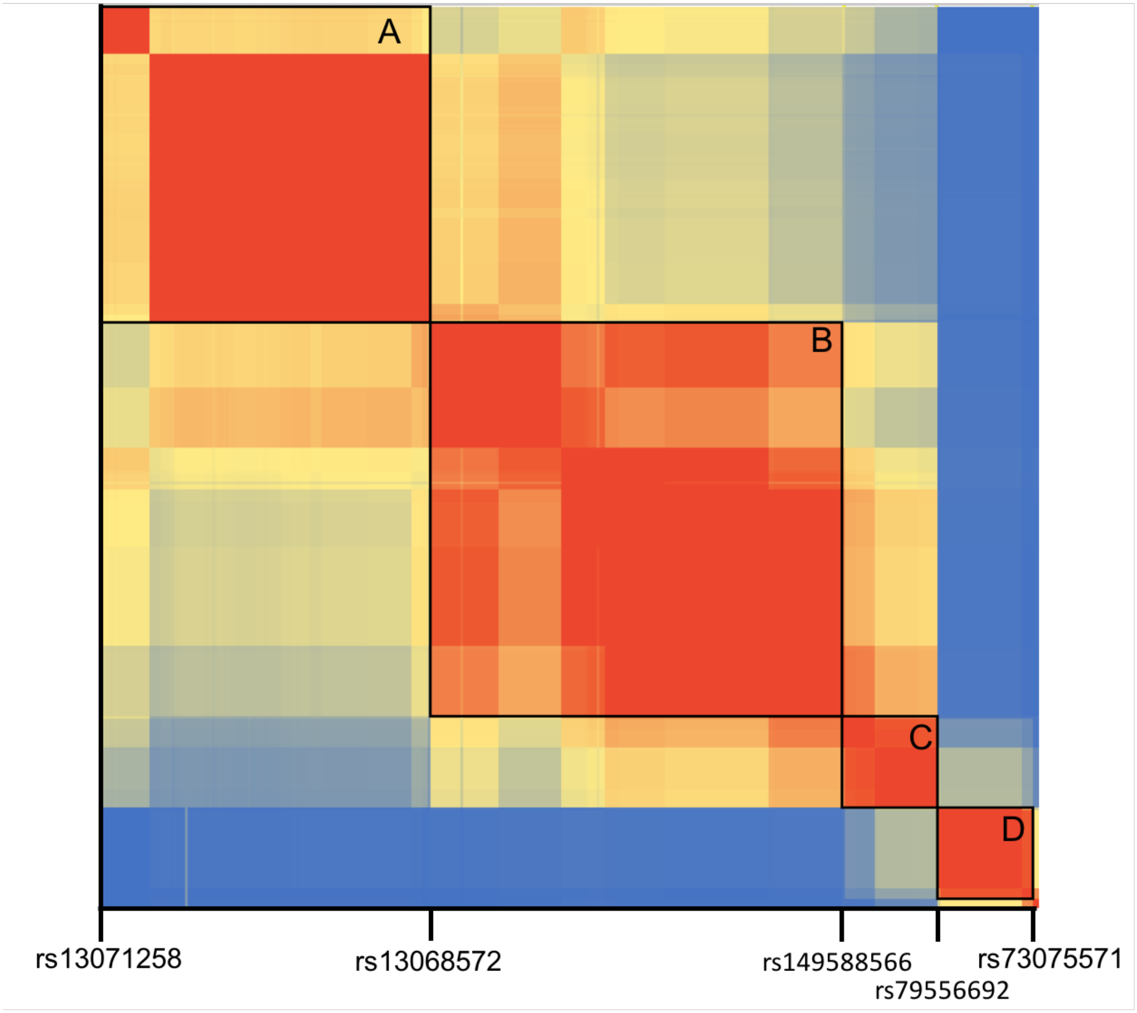


**Supplemental Figure 1.** Linkage Disequilibrium (LD) between SPrime identified introgressed variants within the segment (chr3:45843242-46654616) containing the COVID-19 associated haplotype. This heatmap shows the pairwise $r^2$ LD between each one of the 361 Sprime identified variants (rows) against every other one of the variants (columns). The color scale is a gradient between blue for $r^2 = 0$ and red for $r^2 = 1$. Black boxes indicate the 4 major LD blocks (min r2 = 0.34) with labels A-D denoting our naming scheme for these blocks. Labeled SNPs represent the boundaries between LD blocks.
